## Supplementary figures for "Simultaneous quantification of mRNA and protein in single cells reveals post-transcriptional effects of genetic variation"

Department of Genetics, Cell Biology and Development,
University of Minnesota
Minneapolis, MN, U.S.A.

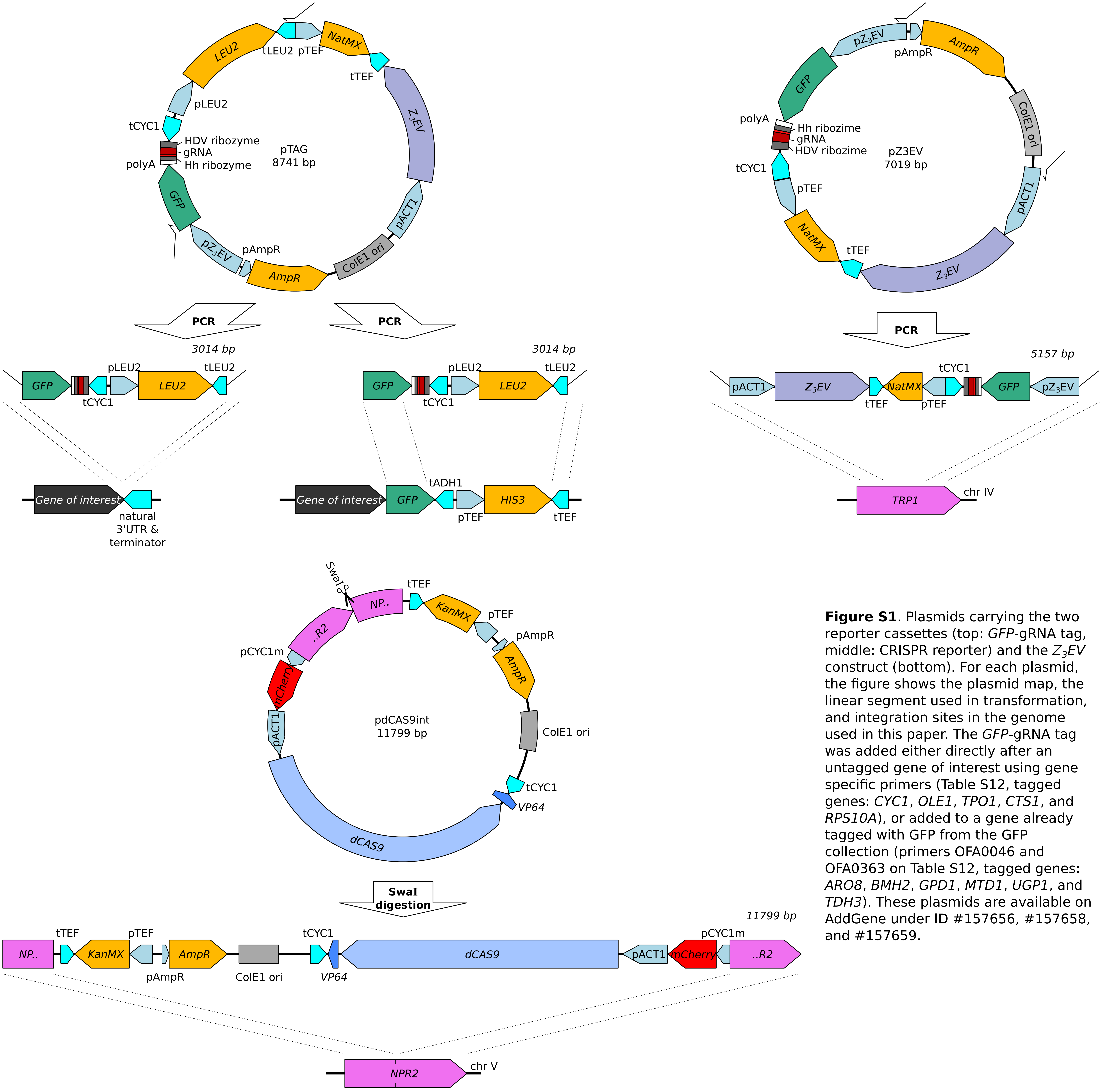


**Figure S1**. Plasmids carrying the two reporter cassettes (top: *GFP*-gRNA tag, middle: CRISPR reporter) and the *Z₃EV* construct (bottom). For each plasmid, the figure shows the plasmid map, the linear segment used in transformation, and integration sites in the genome used in this paper. The *GFP*-gRNA tag was added either directly after an untagged gene of interest using gene specific primers (Table S12, tagged genes: *CYC1*, *OLE1*, *TPO1*, *CTS1*, and *RPS10A*), or added to a gene already tagged with GFP from the GFP collection (primers OFA0046 and OFA0363 on Table S12, tagged genes: *ARO8*, *BMH2*, *GPD1*, *MTD1*, *UGP1*, and *TDH3*). These plasmids are available on AddGene under ID #157656, #157658, and #157659.

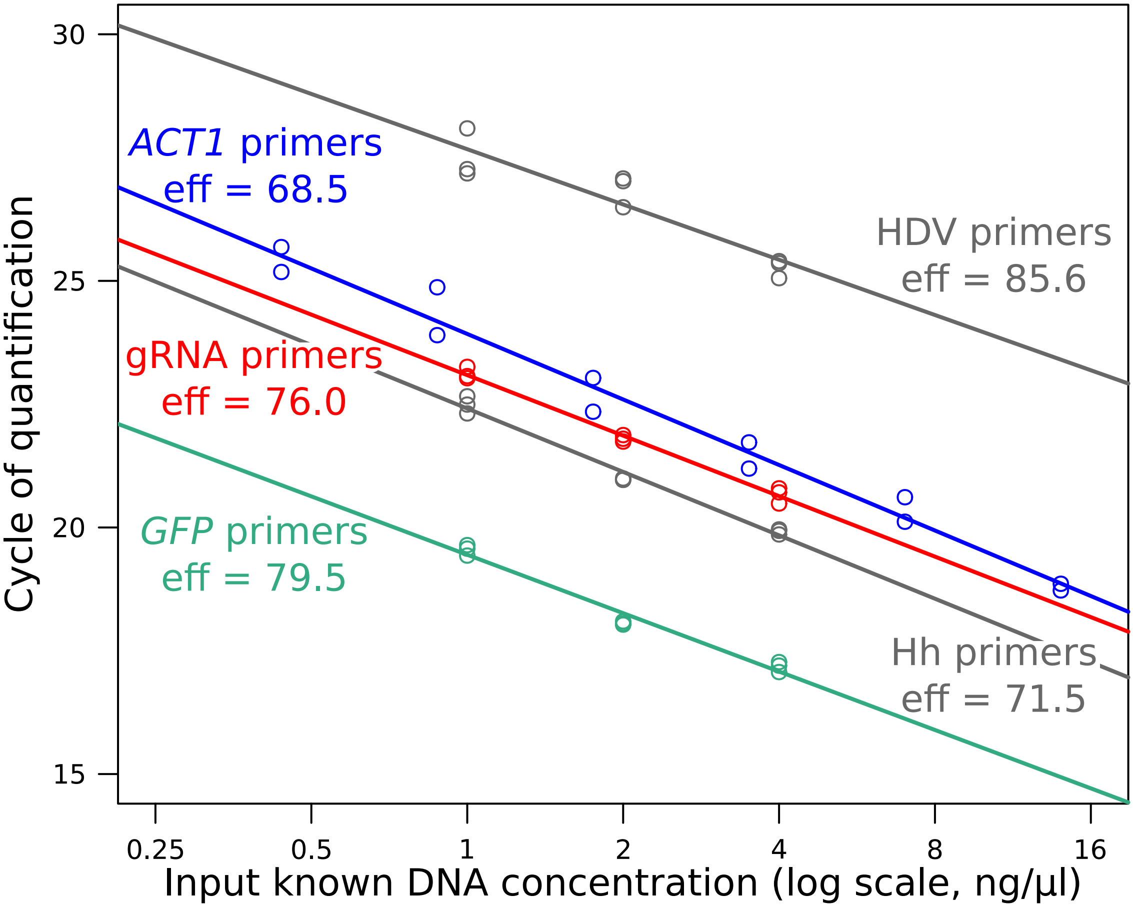


**Figure S2**. qPCR calibration plot for the four primer pairs, using a range of known DNA concentrations. The linear regression coefficients (shown as solid lines) were used to estimate RNA abundance during RT-qPCR for Figures 1, 2 and 6. Efficiency ("eff") was calculated as (2^(-1/slope)-1)*100.

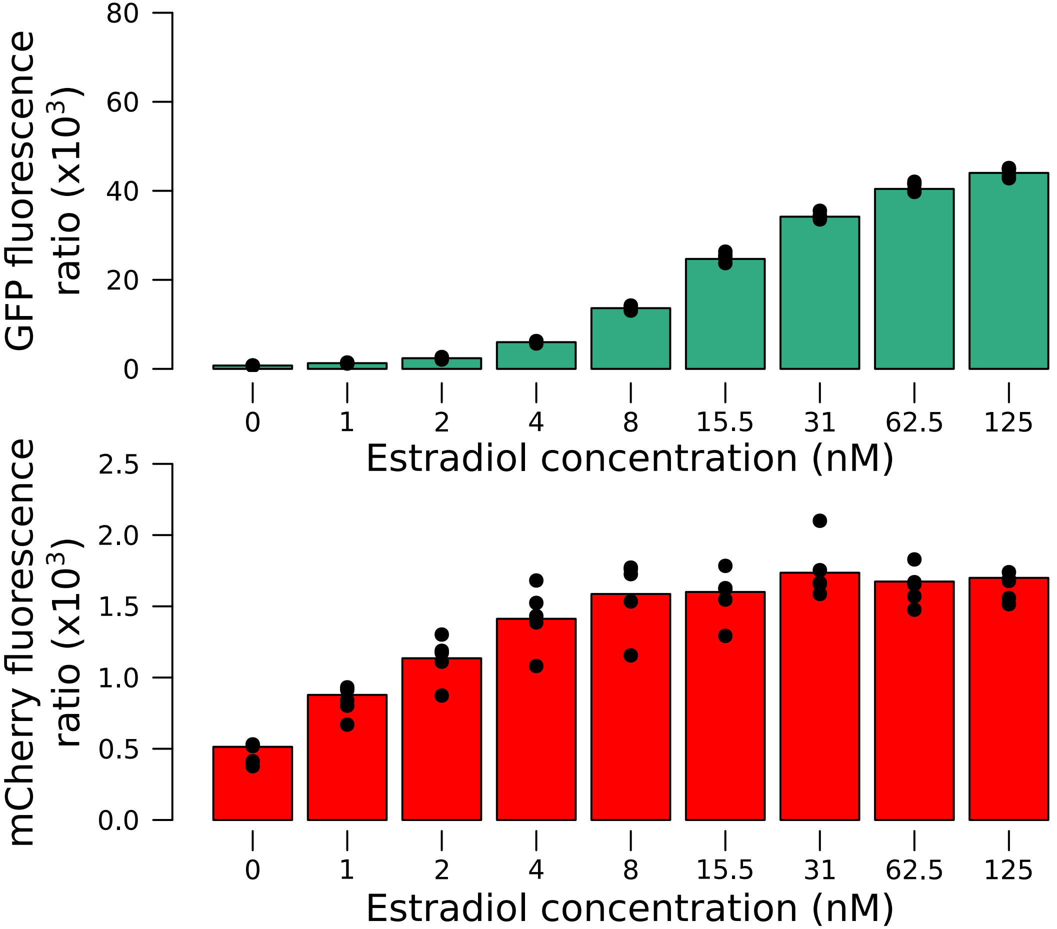


**Figure S3**. Quantitative response of the reporter measured using the $Z_3EV$ system in an RM strain. Increasing concentrations of estradiol drove higher expression of the tagged gene. Values correspond to the fluorescence ratio at the end of the exponential growth phase. Cells were grown in SC medium.

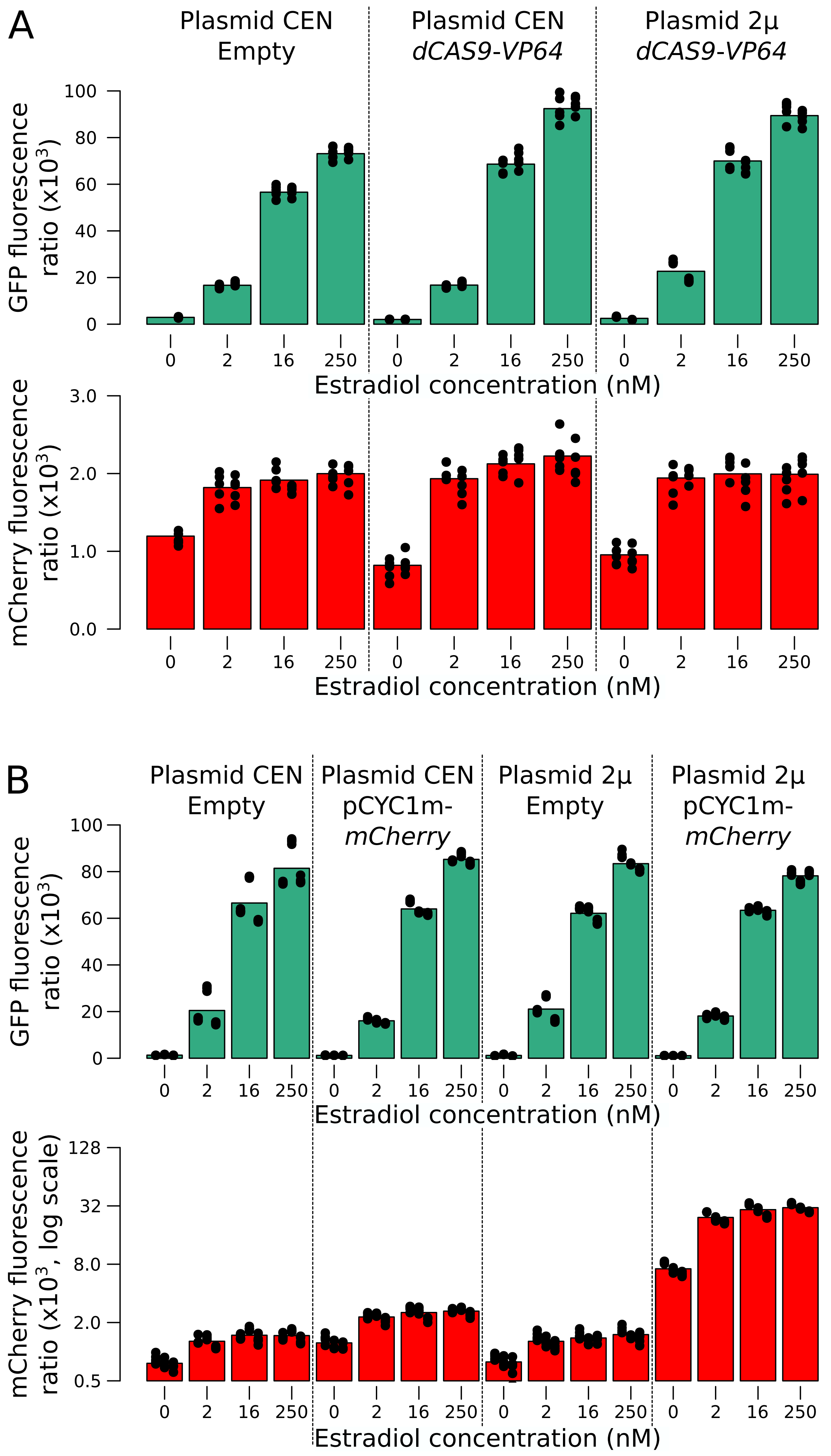


**Figure S4**. *mCherry* and *GFP* expression as a function of additional reporter component copies in a range of estradiol concentrations. (A) Additional copies of the *dCAS9-Vp64* gene using a low-copy number (CEN) or a high-copy number (2µ) plasmid affected neither *GFP* nor *mCherry* expression. Two replicates per condition are shown as stacked columns of points, each with five points that show the values at the time points at which measurements were taken. (B) Additional copies of mCYC1p-*mCherry* increased red fluorescence in all concentrations of estradiol, but did not extend the linear range of *mCherry* expression. Three replicates per condition. Cells were grown in SC medium.

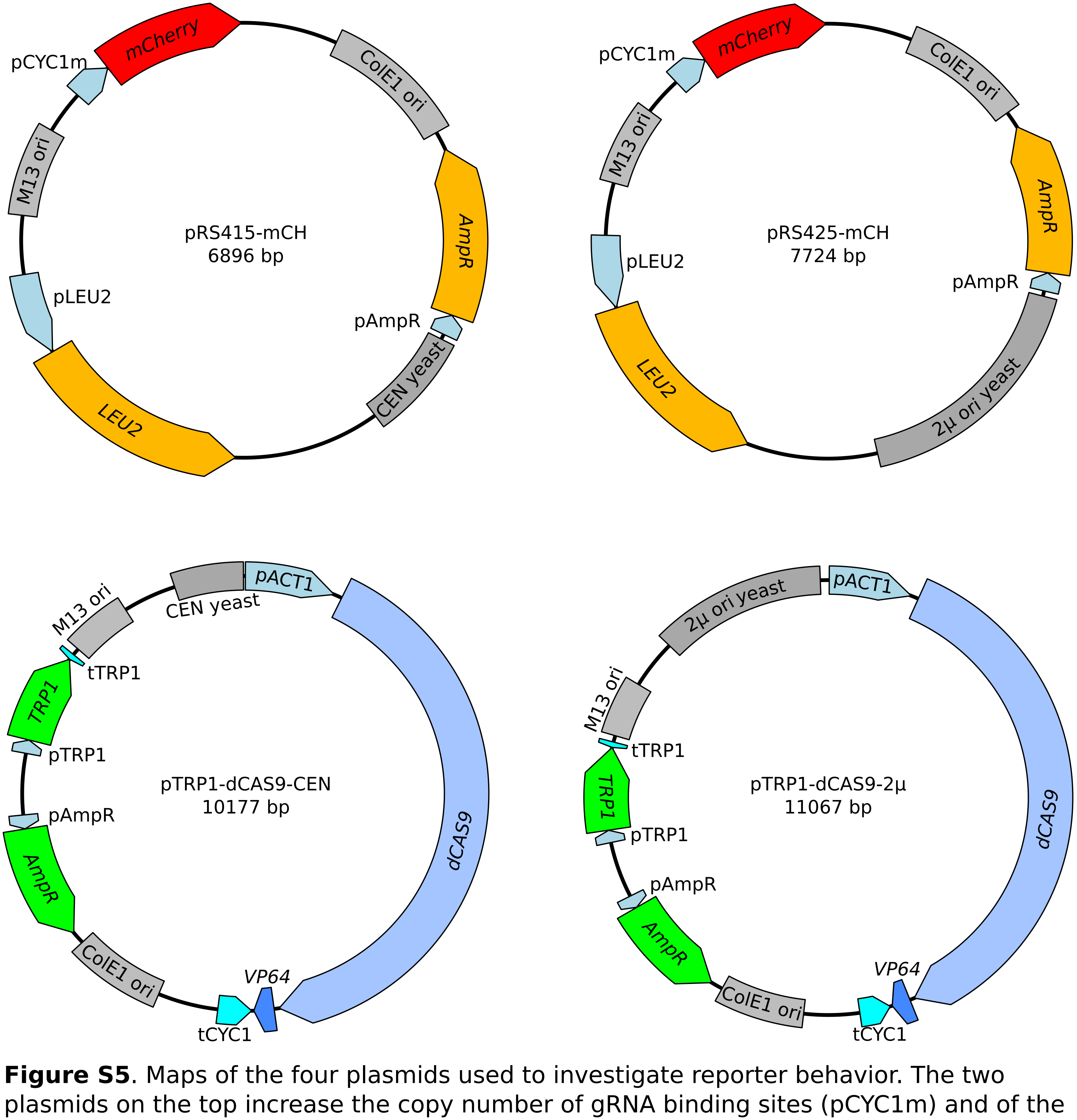


**Figure S5**. Maps of the four plasmids used to investigate reporter behavior. The two plasmids on the top increase the copy number of gRNA binding sites (pCYC1m) and of the *mCherry* gene. The two plasmids on the bottom increase the copy number of *dCAS9-VP64*.

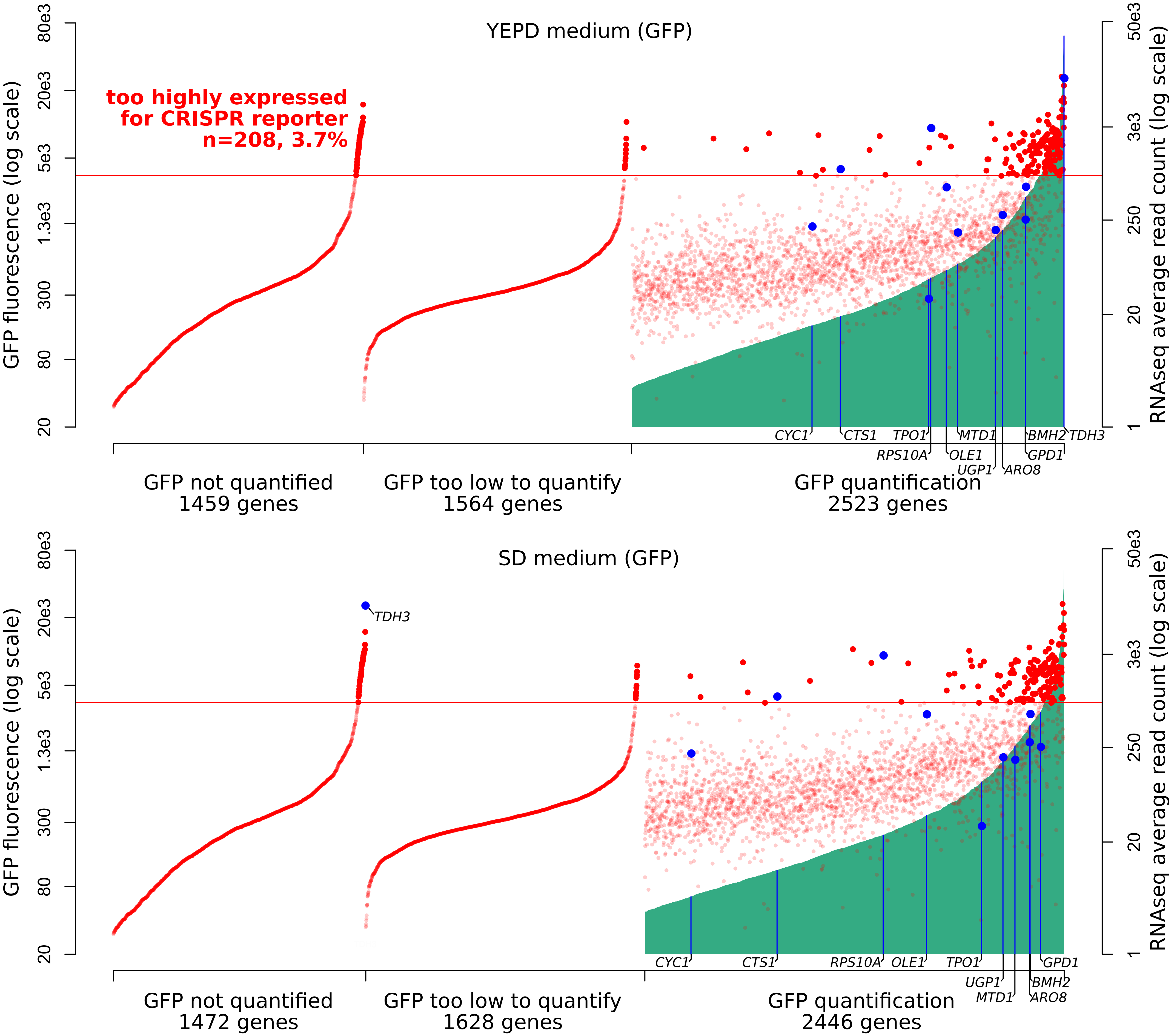


**Figure S6**. Expression ranges of *S. cerevisiae* genes. On the x-axis, 5,546 genes were ordered using two measures of expression. First, genes were ordered on the basis of GFP fluorescence as reported in the GFP collection paper (Huh *et al.* 2003) (green vertical lines). About half of the genes in the genome had available quantifications. Top panel: GFP measures in YEPD medium, bottom: SD medium. Second, genes were ordered on the basis of mRNA abundance in YNB medium as determined by RNA sequencing (red dots, data are averaged across 1,012 BY / RM segregants reported in Albert, Bloom *et al.* 2018). Genes examined in the present study are indicated in blue. The red horizontal line indicates the estimated threshold below which mRNA abundance of a gene can be quantified by the CRISPR reporter.

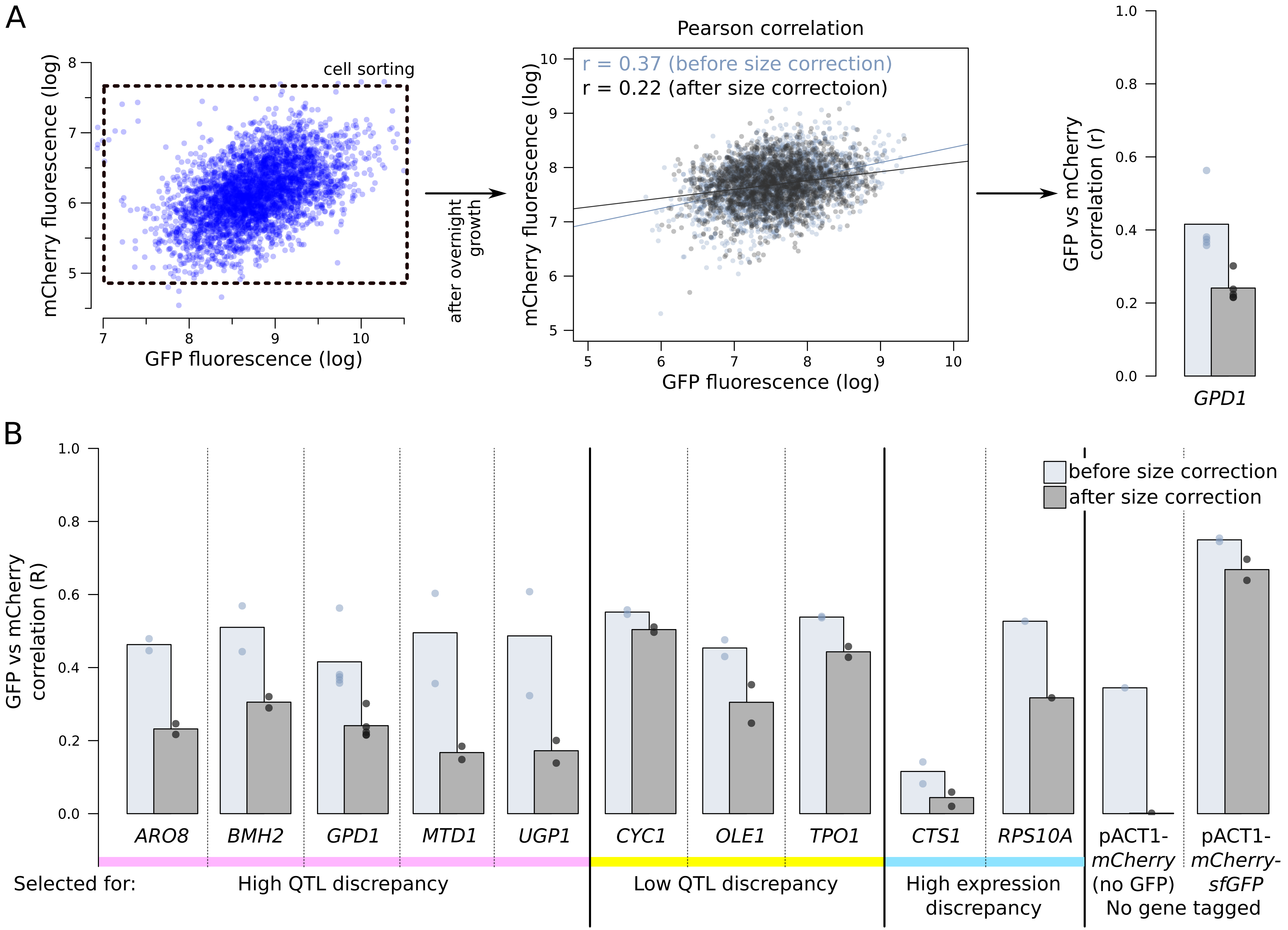


**Figure S7**. mCherry and GFP fluorescence correlations. (A) Following overnight growth after sorting, segregant populations were analysed by flow cytometry. We calculated mRNA and protein abundance correlations (Pearson's r) with and without correction for cell size (see Figure S8). (B) Genes with higher reported discrepancies between mRNA-QTLs and protein-QTLs tended to show lower correlation between mRNA (mCherry) and protein (GFP) abundance than genes with more similar prior QTL results. We used two controls. First, in a segregant population with no gene tagged and with mCherry under the control of a constitutive *ACT1* promoter (see Figure S10), we expected no mCherry / GFP correlation. Second, a segregant population expressing a protein fusion of *mCherry* and *GFP* under control of the *ACT1* promoter provided an upper bound for the possible mCherry / GFP correlation.

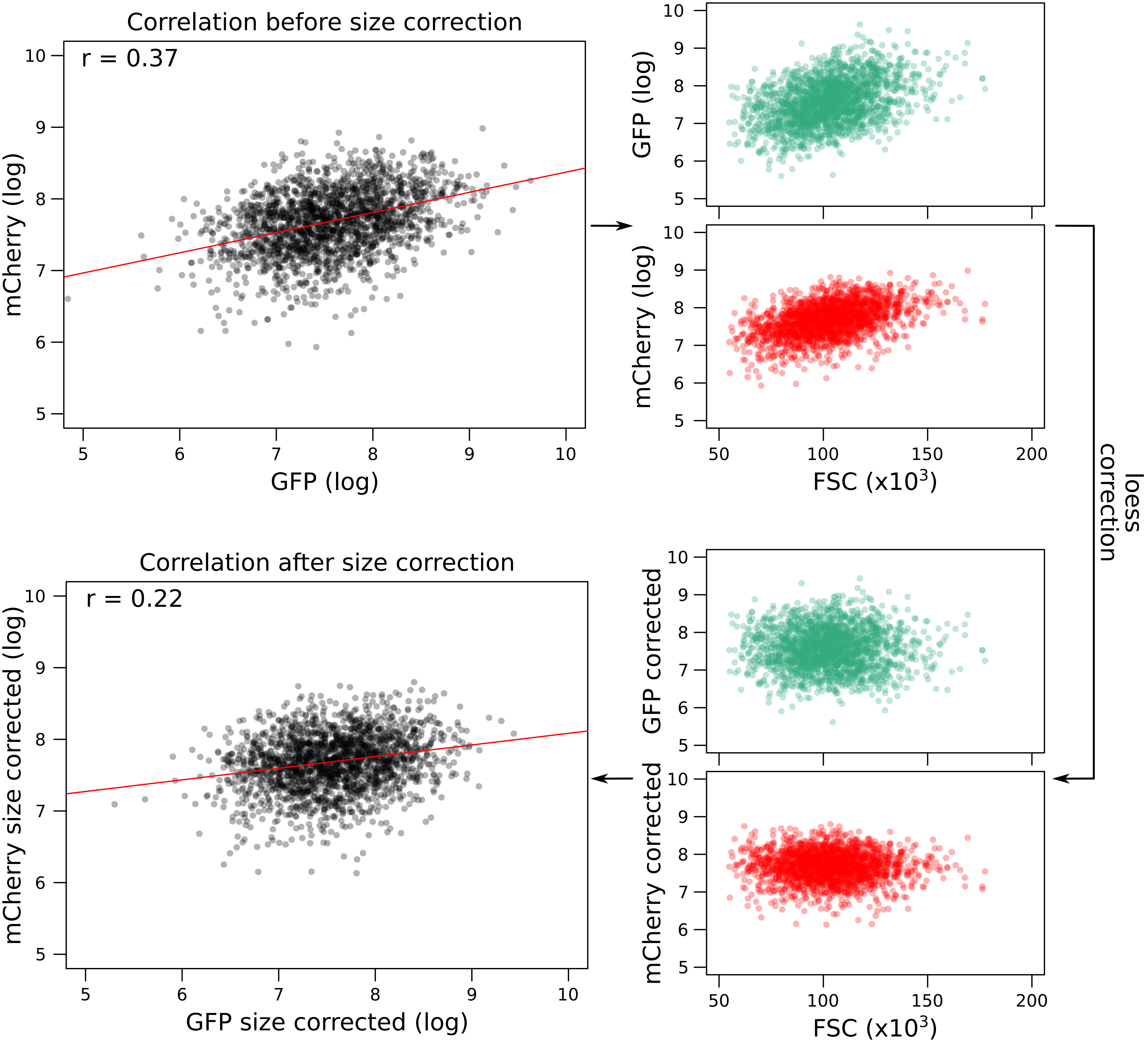


**Figure S8**. Correction of fluorescence measures for cell size to prevent spurious correlations between mCherry and GFP due to shared correlations with cell size. As an example, the figure shows data from *GPD1-GFP*-gRNA.

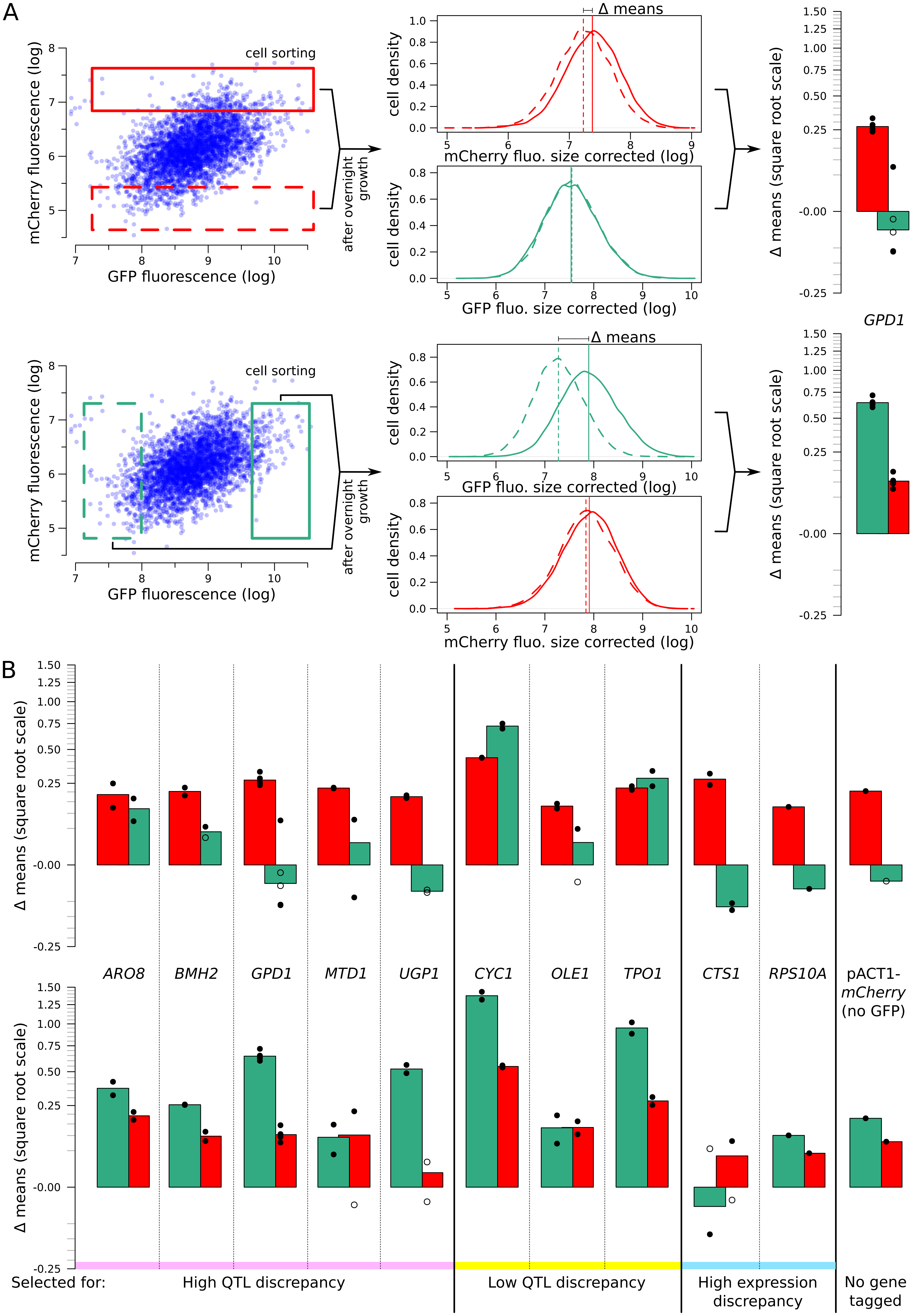


**Figure S9**. Heritability of mCherry and GFP fluorescence. (A) Flow cytometry measures of high and low populations after overnight growth. The difference of fluorescence means (Δ means) between the high and low populations that is retained after multiple generations of growth reflects how much of the variation among single cells is due to genetic variation. When measuring the same fluorophore used to collect extreme cells, Δ means reflects the heritability of that fluorophore. When measuring the other fluorophore than that used in sorting (e.g., measuring GFP in a pair of populations that had been sorted based on different mCherry levels), Δ means reflects the genetic correlation between mRNA and protein. (B) Significant heritability was observed for all fluorescent measures across genes, except for GFP in *CTS1*. Genes with higher reported discrepancies between mRNA-QTLs and protein-QTLs tended to have lower genetic correlations between mRNA and protein than genes with more concordant prior QTL results. Bar plots of Δ means are square-root transformed to visually emphasize smaller values. Empty dots correspond to non-significant differences. Filled dots indicate significant differences at a t-test p-value threshold of $10^{-5}$.

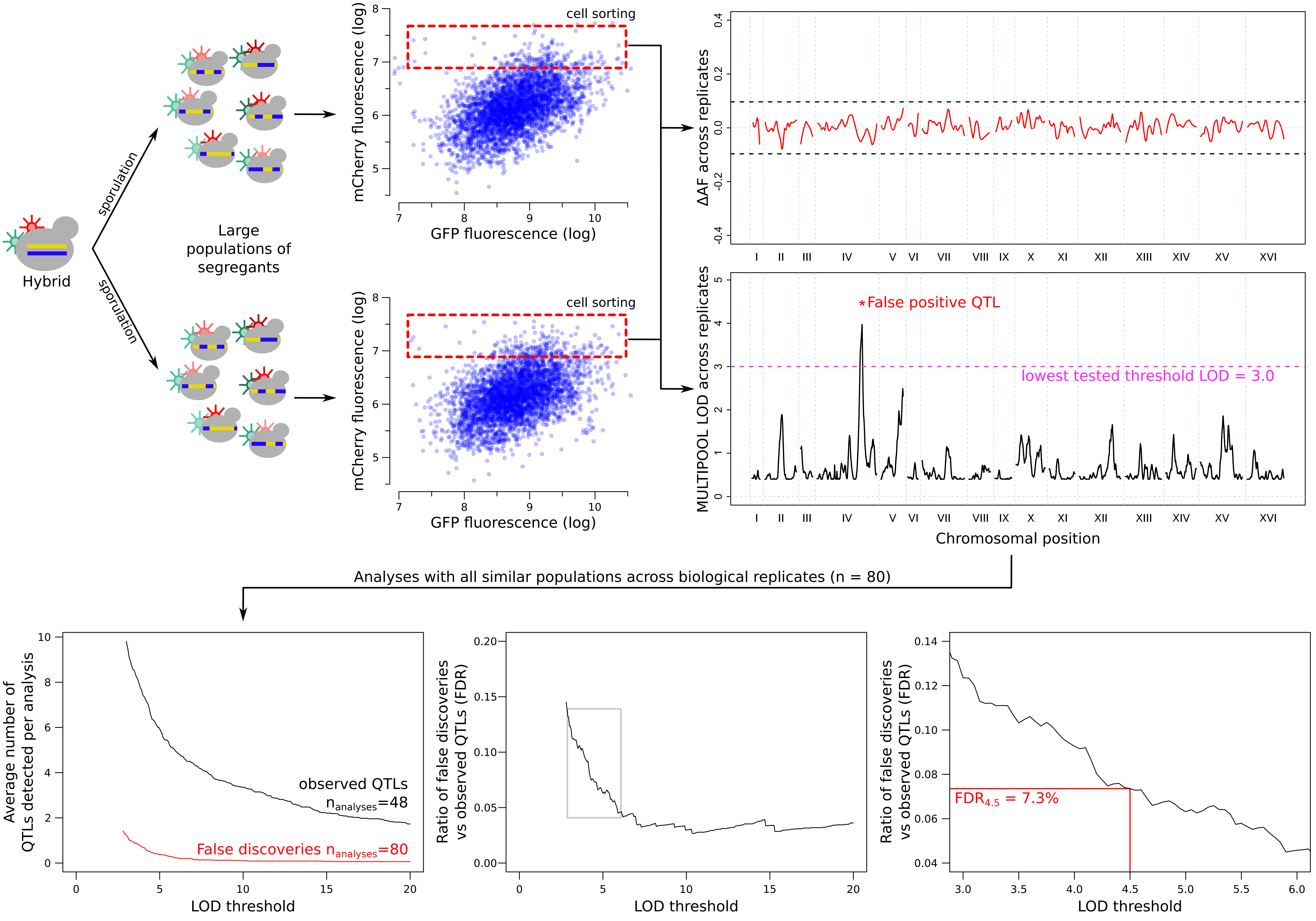


**Figure S10**. Estimation of the false discovery rate (FDR) from comparisons across replicated sort experiments. Allele frequencies were compared between replicate populations collected using the same sort gates. The example shown here is based on data from the *GPD1* gene. Any QTLs between these populations were considered to be false discoveries. Increasing the LOD threshold reduced both the number of observed QTLs (from the high vs low population comparisons) and false QTLs (from the comparison of the same population type across replicates). FDR was calculated as a function of the LOD threshold (*thr*): $FDR_{thr} = (NrepQTL_{thr} / Nrep) / (NfluoQTL_{thr} / Nfluo)$, where $NrepQTL_{thr}$ is the number of QTLs that exceeded a LOD threshold of *thr* (false discoveries), *Nrep* is the number of inter-replicate comparisons (*Nrep* = 80), $NfluoQTL_{thr}$ is the number of significant fluorescence QTLs (GFP or mCherry) at a LOD threshold of *thr*, and *Nfluo* is the number of high-low fluorescence comparisons (*Nfluo* = 48; excluding the control experiment in which no gene was tagged; Figure S11). The threshold of LOD = 4.5, which corresponded to an FDR of 7.3%, is indicated in the figure.

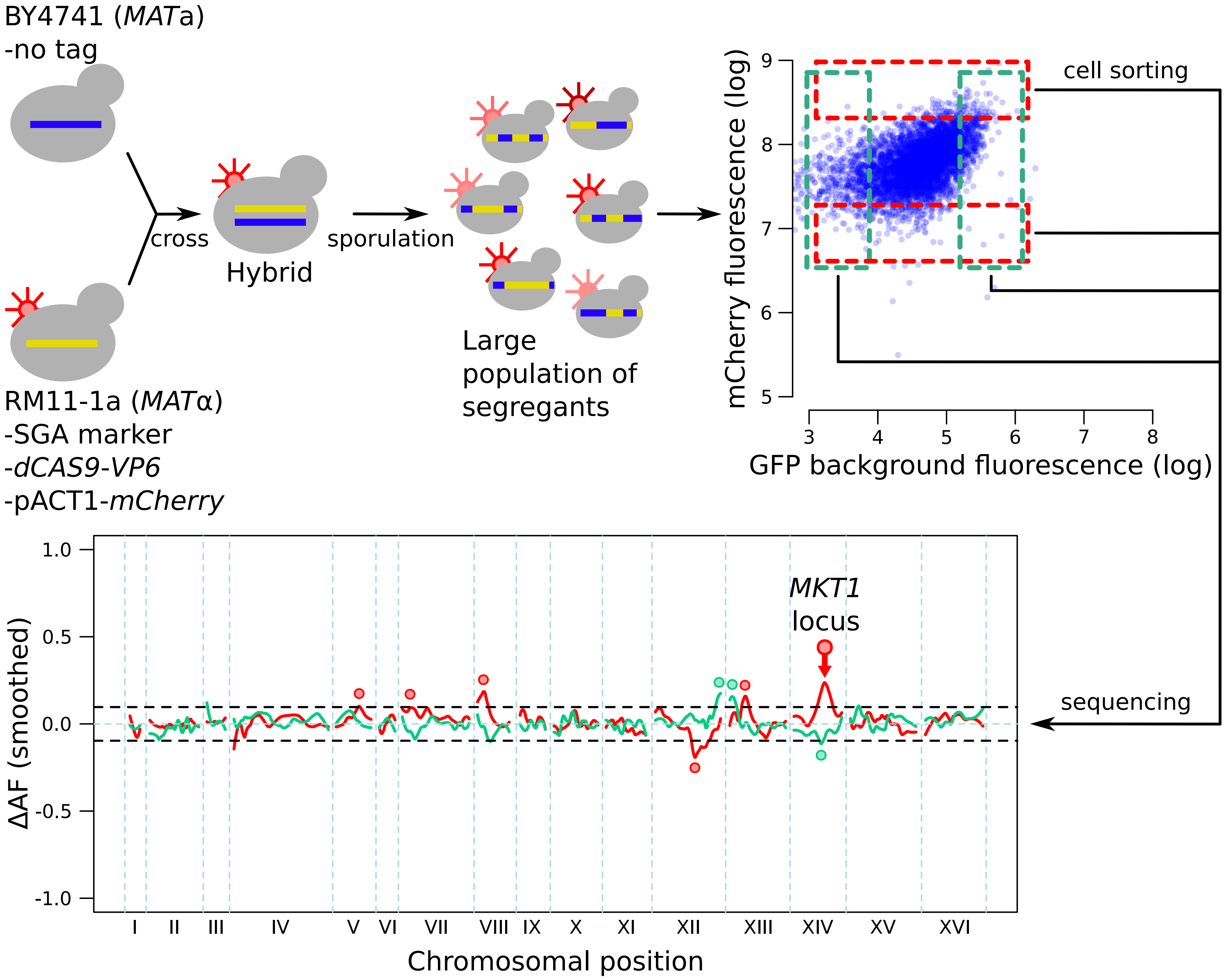


**Figure S11**. Control mapping experiment, in which no gene was tagged with GFP or the gRNA, and in which the mCherry gene was under control of a constitutively active *ACT1* promoter sequence. The highlighted mCherry QTL on chromosome XIV, which overlaps the *MKT1* locus, was also present in all experiments with tagged genes (Figure 4), where it always had the same direction of effect as in this control experiment. While the highly pleiotropic *MKT1* locus (Albert, Bloom *et al.* 2018) could truly influence all ten genes we studied, we cannot rule out that this locus may affect mCherry fluorescence independently of the gRNA. We therefore excluded QTLs from tagged genes in this region from our analysis. We did not exclude the remaining mCherry QTLs visible in the figure because these loci were not uniformly present for tagged genes. These loci could represent trans-acting influences on the *ACT1* promoter sequence that drives mCherry expression in this control experiment, but not in experiments with tagged genes under the control of their native promoters. The control experiment also identified three loci that affected GFP background fluorescence, likely by altering cellular abundance of autofluorescent compounds such as NAD, aromatic amino acids, or flavins. The intensity of this autofluorescence was much lower than the GFP signal for the genes we studied (see for example Figures S9 and S10), such that in those experiments, the signal from the *GFP* gene tags dominates over background fluorescence.

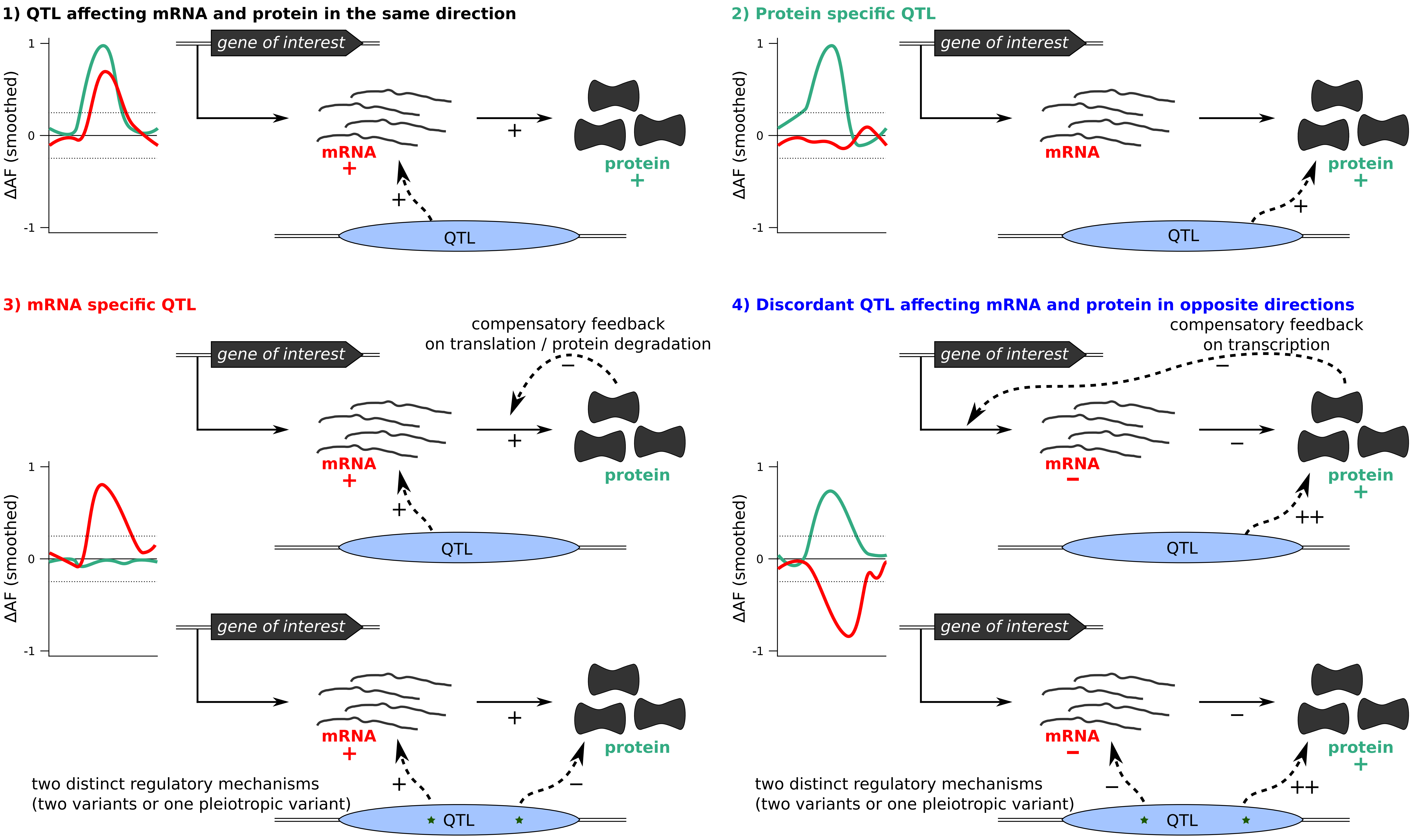


**Figure S12**. Classification of QTLs into four groups based on their effect on protein and / or mRNA. Differences in allele frequency (ΔAF) are illustrations rather than observed data. Examples of possible underlying mechanisms for each scenario are shown.

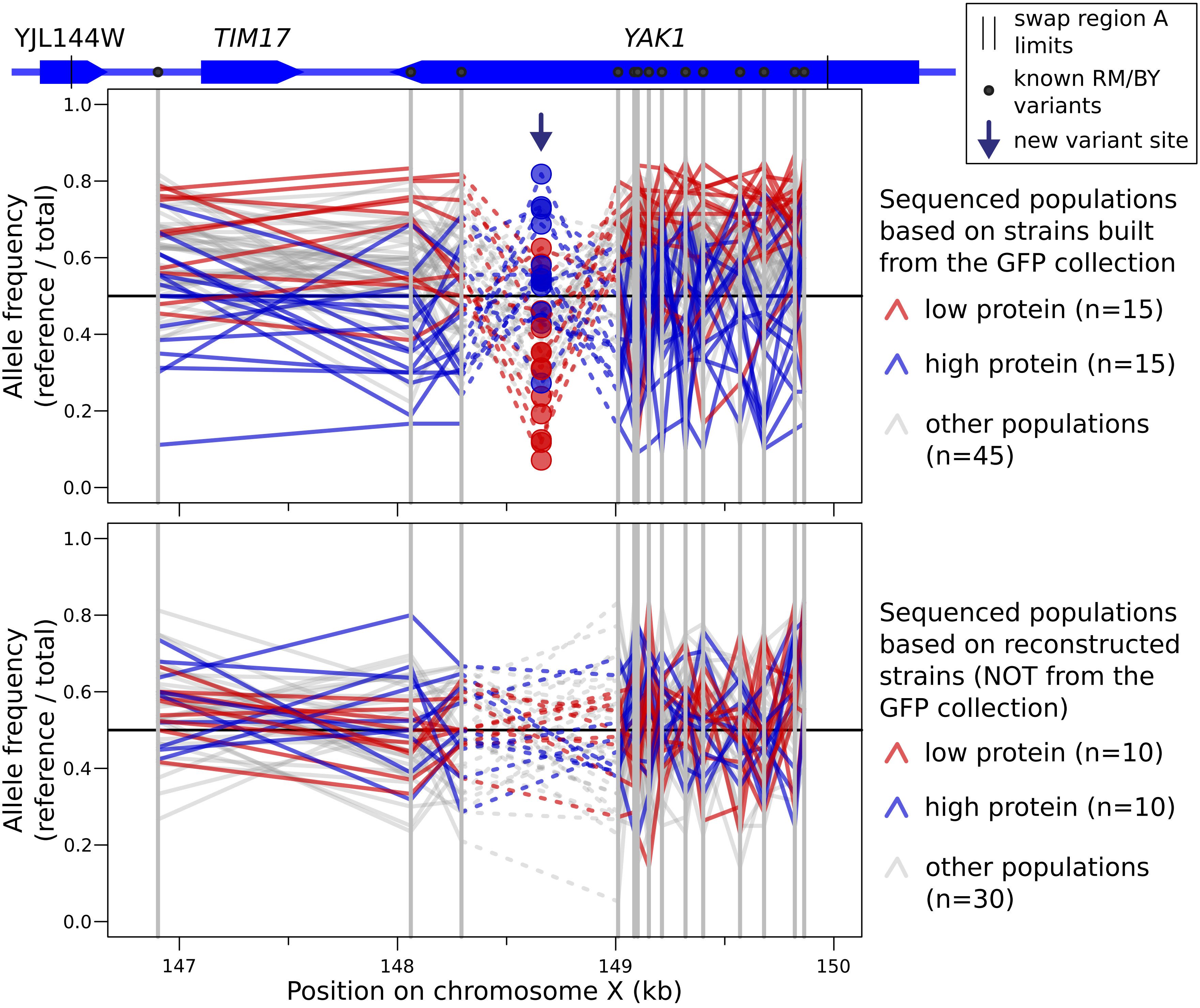


**Figure S13**. Allele frequency of variants in tile A for every sequenced segregant population, separated between populations based on BY strains from the GFP collection (top), or on a separate BY strain into which we had engineered the GFP tag (bottom). For each sequenced population, a line traces the allele frequency of neighboring variants along the genome. Known BY / RM variants are indicated by solid grey vertical lines. The new A/G single nucleotide variant at 148,659 bp is indicated by an arrow. This mutation was detected as a heterozygous site in the majority of cell populations constructed from GFP collection strains (72 out of 75 populations). The mutation was detected in no populations constructed from strains that did not directly originate from the GFP collection (0 out of 50 populations). The apparent allele frequency inversion at this variant compared to nearby variants is due to the fact that the alternative “G” allele at this variant is carried by a derivative of the reference BY strain.

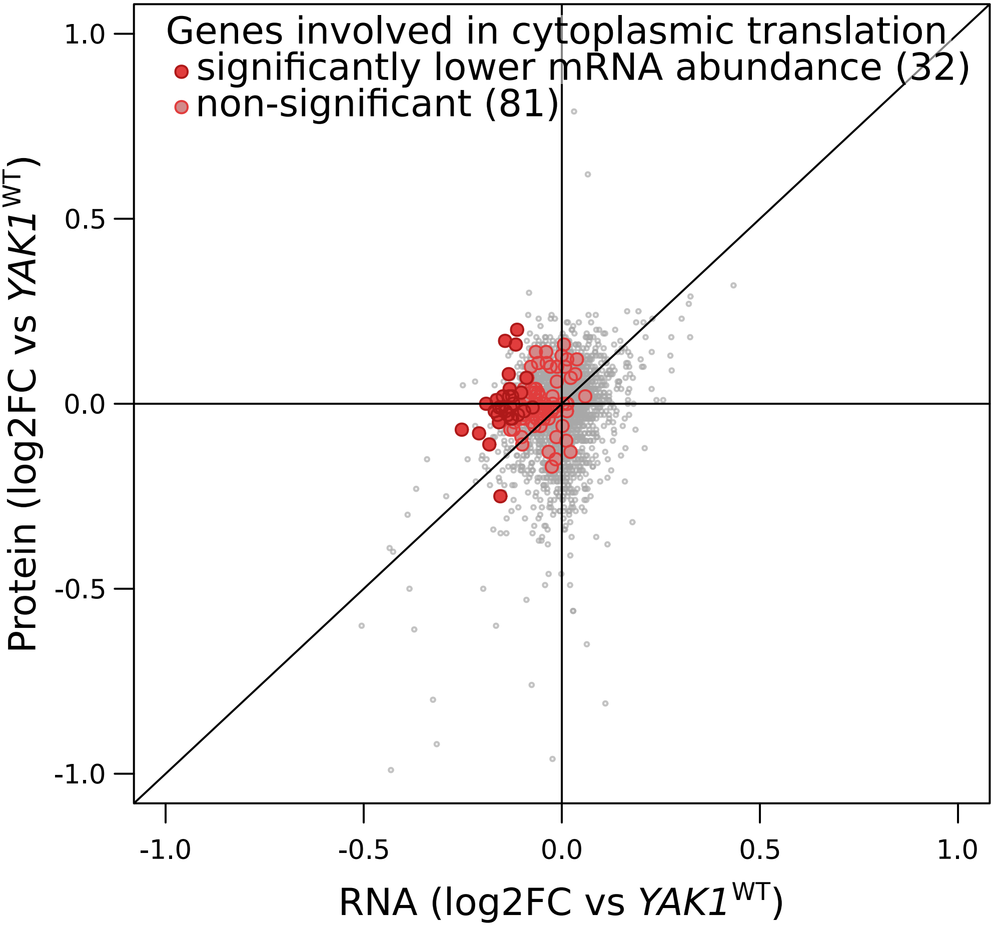


**Figure S14**. *YAK1*$^{Q578*}$ effect on the expression of genes involved in cytoplasmic translation. Each point corresponds to a gene for which mRNA levels (x-axis) and protein levels (y-axis) were quantified by RNA sequencing and mass spectrometry, respectively. Genes with lower mRNA abundance were significantly enriched in cytoplasmic translation: 32 genes annotated to this functional category were significant (dark red; Benjamini-Hochberg adjusted p-value, < 0.05), compared to 81 that were not significant (light red): hypergeometric test p-value = $2.2\times10^{-14}$. log2FC: $\log_2$ of fold-change.
